# Glucose-6-phosphate dehydrogenase deficiency accelerates pancreatic acinar-to-ductal metaplasia

**DOI:** 10.1101/2023.11.06.565895

**Authors:** Megan D. Radyk, Barbara S. Nelson, Christopher J. Halbrook, Alexander Wood, Brooke Lavoie, Lucie Salvatore, Gabriel Corfas, Justin A. Colacino, Yatrik M. Shah, Howard C. Crawford, Costas A. Lyssiotis

## Abstract

Activating mutations in *KRAS* extensively reprogram cellular metabolism to support the continuous growth, proliferation, and survival of pancreatic tumors. Targeting these metabolic dependencies are promising approaches for the treatment of established tumors. However, metabolic reprogramming is required early during tumorigenesis to provide transformed cells selective advantage towards malignancy. Acinar cells can give rise to pancreatic tumors through acinar-to-ductal metaplasia (ADM). Dysregulation of pathways that maintain acinar homeostasis accelerate tumorigenesis. During ADM, acinar cells transdifferentiate to duct-like cells, a process driven by oncogenic *KRAS*. The metabolic reprogramming that is required for the transdifferentiation in ADM is unclear. We performed transcriptomic analysis on mouse acinar cells undergoing ADM and found metabolic programs are globally enhanced, consistent with the transition of a specialized cell to a less differentiated phenotype with proliferative potential. Indeed, we and others have demonstrated how inhibiting metabolic pathways necessary for ADM can prevent transdifferentiation and tumorigenesis. Here, we also find NRF2-target genes are differentially expressed during ADM. Among these, we focused on the increase in the gene coding for NADPH-producing enzyme, Glucose-6-phosphate dehydrogenase (G6PD). Using established mouse models of *Kras^G12D^*-driven pancreatic tumorigenesis and G6PD-deficiency, we find that mutant *G6pd* accelerates ADM and pancreatic intraepithelial neoplasia. Acceleration of cancer initiation with G6PD-deficiency is dependent on its NADPH-generating function in reactive oxygen species (ROS) management, as opposed to other outputs of the pentose phosphate pathway. Together, this work provides new insights into the function of metabolic pathways during early tumorigenesis.

## INTRODUCTION

Most patients with pancreatic ductal adenocarcinoma (PDAC) are diagnosed late in progression, often with metastasized disease^1^. PDAC has a low 5-year survival rate of 12%, due in part to a lack of early detection methods and effective therapies^2^. Oncogenic *KRAS* mutations are initiating events in PDAC and present in over 90% of pancreatic cancers^3^. Tumorigenesis in PDAC is hypothesized to progress stepwise beginning with acinar-to-ductal metaplasia (ADM), followed by pancreatic intraepithelial neoplasia (PanIN), leading to invasive carcinoma. Lineage tracing studies in PDAC mouse models demonstrates that mutant *Kras*-expressing acinar cells can undergo ADM and give rise to cancer^4, 5^. ADM is a reversible, wound healing response after pancreatic injury or inflammation. In ADM, acinar cells transdifferentiate to ductal progenitor-like cells, proliferate, and reconstitute lost/damaged tissue^6–8^. Upon healing, ductal progenitor cells redifferentiate to acinar cells and resume normal acinar function. However, oncogenic *Kras* mutations block redifferentiation, which leads to persistent ADM and permits progression to neoplastic lesions and PDAC. ADM can be blocked by inhibiting signaling pathways necessary for transdifferentiation^9–12^. In addition, targeting pathways that drive tumorigenesis, such as oncogenic *Kras*, can revert ADM and PanIN lesions back to normal acinar cells^9, 10^.

Pancreatic cancer cells require continuous supply of biosynthetic precursors and energy to sustain growth and proliferation^13–15^. To meet this demand, cells undergo metabolic reprogramming to promote aerobic glycolysis (the Warburg Effect) where glucose is diverted into anabolic pathways to generate biosynthetic precursors and energy^16^. Pancreatic cancer cells also utilize noncanonical glutamine metabolism and alter their redox and antioxidant pathways^17, 18^. Additionally, oncogenic *KRAS* selectively activates the non-oxidative branch of the pentose phosphate pathway (PPP) through both enhanced glycolysis leading to overflow metabolism and transcriptional activation of *Rpia* and *Rpe*^16, 17, 19^. This tilts the production of PPP toward the production of 5-carbon sugars independent of NADPH from the oxidative branch. In contrast, the oxidative PPP produces ribose 5-phosphate and is the major source of NADPH in the cell^16, 19^. Metabolic reprogramming in PDAC has been well characterized, but there is a paucity of data on how metabolism drives precancerous ADMs and PanINs. However, it has been reported that cancer initiation and progression in PDAC relies on tight reactive oxygen species (ROS) control^20–22^.

Glucose-6-phosphate dehydrogenase (G6PD) is the first and rate-limiting enzyme in the oxidative PPP. G6PD also produces NADPH via the conversion of glucose 6-phosphate to 6-phosphogluconolactone. NADPH can be used for biosynthetic pathways (fatty acids, cholesterol, deoxyribonucleotides) and cellular antioxidant systems (glutathione and thioredoxin) that mitigate ROS^23^. G6PD-deficiency is an X-linked disorder and the most common enzyme defect in humans^24^. Red blood cells are highly sensitive to decreased oxidative PPP flux, as they possess limited metabolites for the generation of NADPH via alternate pathways. Thus, G6PD-deficiency presents as chronic or hemolytic anemia and includes 186 known mutations^25^. Retrospective studies show that patients with G6PD-deficiency may be protected from stomach, colon, and liver cancers, but not others^26, 27^. Recently, it was shown that loss of TIGAR, a protein that supports activation of the oxidative PPP, delays PanIN lesions, but enhances metastasis^22^. Thus, there is complex regulation of the oxidative PPP and ROS during cancer initiation, development, and progression.

Here we use established mouse models of *Kras^G12D^*-driven pancreatic tumorigenesis to determine which metabolic pathways are important in the development of precancerous lesions. We find that decreased oxidative PPP flux, by G6PD-deficiency, accelerates ADM and PanIN formation via increased ROS. This metabolic pathway and others identified from RNA-sequencing (RNA-seq) provide insight into the metabolic changes during cancer initiation and may inform future therapies and risk factors.

## RESULTS

### Metabolic changes during acinar-to-ductal metaplasia

Oncogenic *KRAS* regulates metabolic reprogramming in PDAC cells^16, 17^ and is required for the initiation of pancreatic cancer^9^. To investigate *Kras^G12D-^*driven changes that occur during pancreatic cancer initiation, we performed transcriptomics on *ex vivo* cultured acinar cells undergoing acinar-to-ductal metaplasia (ADM). Mutant *Kras* elicits an inflammatory response in pancreatic tissue^28^; thus, we activated oncogenic *Kras ex vivo* as to identify ADM cell autonomous *Kras^G12D^*-driven changes and control for differences in inflammation between mutant *Kras*-expressing and wild-type pancreas. We isolated acinar cells from *LSL-Kras^G12D/+^*mice and infected the cells with control adenovirus (ad-GFP) or adenovirus expressing Cre recombinase (ad-CRE) to induce expression of mutant Kras^G12D^ (**Figure 1A**). RNA was isolated 24 hours after plating ad-GFP infected acinar cells (GFP) and 24, 48, and 72 hours after plating ad-CRE infected cells (Cre 1/2/3, respectively). GFP cells were only collected 24 hours after plating because acinar cells quickly die without a signal to transdifferentiate (like oncogenic *Kras*). A multidimensional scaling dimension reduction plot based on the RNA-sequencing data shows that samples separate by timepoint on the first dimension, with GFP and Cre1 cells clustering together, and Cre2 and Cre3 cells separating in a manner which reflects the experimental temporality (**Figure 1B**). On the second dimension, cells clustered relative to the mouse of origin.

**Figure 1.**
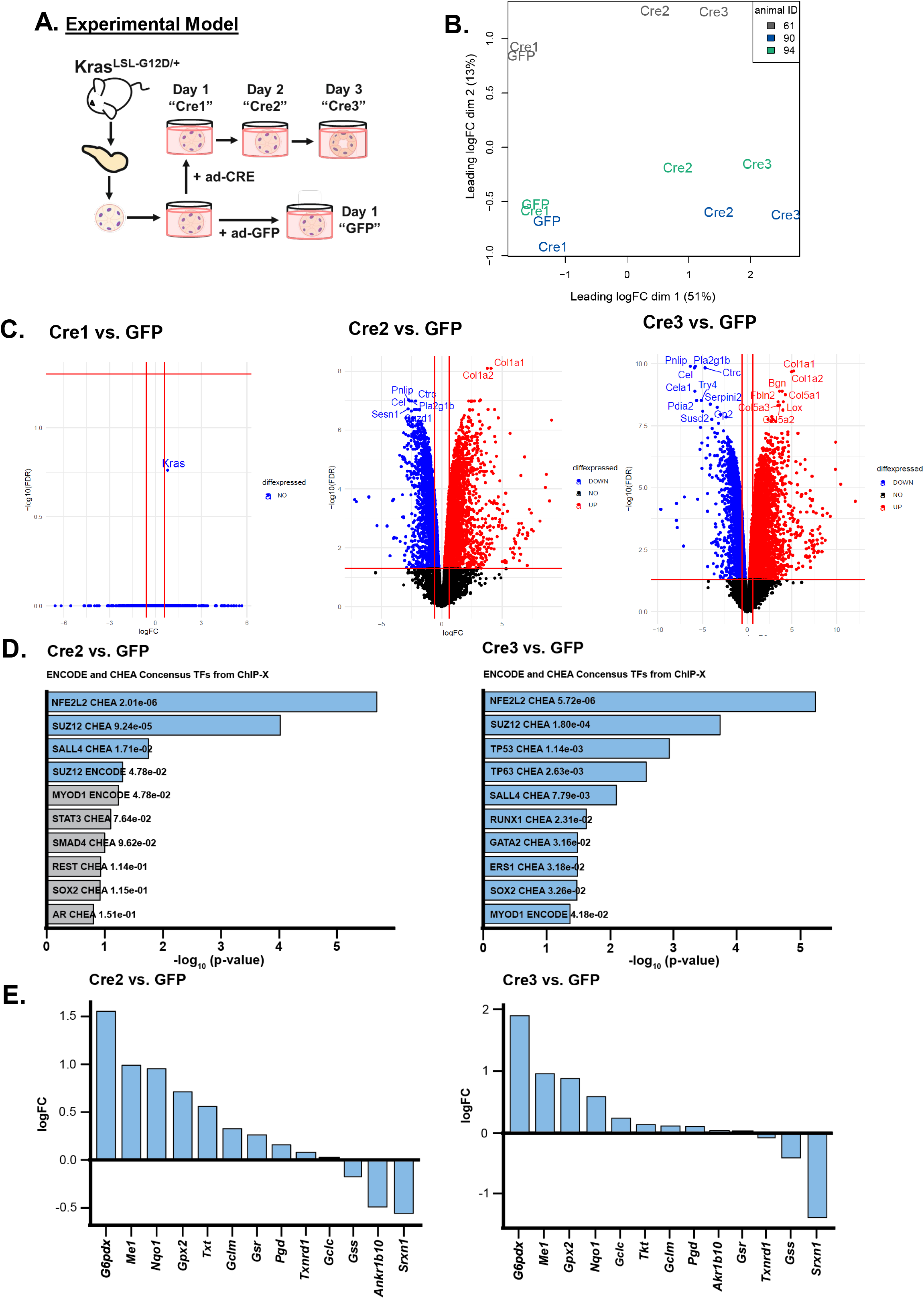
RNA-sequencing shows an NRF2 pathway signature in *ex vivo* acinar cell culture. **A.** Scheme of the experimental model. Pancreata were dissected from *LSL-Kras^G12D/+^* mice and acinar clusters were isolated for *ex vivo* culture. Acinar clusters were grown free-floating in media and were treated with ad-GFP (control) or ad-CRE adenoviruses (to induce mutant *Kras*). Cells were collected after 1, 2, or 3 days in culture and RNA was isolated. At 3 days in culture, acinar cells take on ductal cystic morphology, resembling acinar-to-ductal metaplasia. **B.** Dimensional reduction plot of GFP, Cre1, Cre2, and Cre3 samples from RNA-sequencing. Each color represents one mouse. Left, Cre1 vs. GFP; Center, Cre2 vs. GFP; Right, Cre3 vs. GFP. **C.** Volcano plots of differentially expressed genes when comparing Cre1/2/3 to GFP samples. Upregulated genes are red, downregulated genes are blue. **D.** Consensus transcription factors identified from ENCODE and ChEA databases after inputting the top 500 upregulated genes from RNA-sequencing in the Enrichr platform. Left, Cre2 vs. GFP; Right: Cre3 vs. GFP. Blue bars represent significant p-values, Grey bars represent nonsignificant p-values. **E.** LogFC of several NRF2-target genes in RNA-sequencing dataset. Left: Cre2 vs. GFP; Right: Cre3 vs. GFP.

One day after Cre induction (Cre1), there was no statistically significant change in gene expression compared to GFP; however, the gene with the greatest difference following induction was *Kras* (**Figure 1C**). Two and three days after Cre induction (Cre2, Cre3) there was a dramatic increase differentially expressed genes (**Figure 1C**), including expectedly decreased acinar genes (e.g. *Pnlip* and *Prss1*) and increased ductal or ADM-related genes (e.g. *Onecut1*, *Onecut2*, *Sox9*) (**Supplemental Figure 1**). Gene Set Enrichment Analysis (GSEA)^29, 30^ demonstrated signatures known to be enhanced during ADM, which are present in our transcriptome data from GFP compared to Cre2 acinar cells—prior to ductal formation (**Supplemental Figure 2A**). Genes that are upregulated by KRAS signaling are enriched in *Kras^G12D^*-expressing acinar cells, whereas genes downregulated by KRAS signaling are enriched in control cells. In addition, epithelial-to-mesenchymal transition (EMT), hypoxia, p53 activity, and inflammation signatures are consistent with previous studies showing these pathways are involved in ADM^31–36^. GSEA also revealed metabolic signatures, including fatty acid metabolism, glycolysis, ROS pathways, oxidative phosphorylation, mTORC1 signaling, and cholesterol homeostasis are highly enriched in *Kras^G12D^*-expressing acinar cells (**Supplemental Figure 2B**). Our transcriptomics data is consistent with studies demonstrating elevations in cholesterol metabolism, mTORC1, ROS, and antioxidant programs during ADM^20–22, 37, 38^. However, it is not clear how these pathways are regulated, and whether these metabolic pathways play a role in ADM formation.

### NRF2 pathway signature in Kras-driven acinar-to-ductal metaplasia

To identify transcription factors regulating the metabolic changes observed in ADM, we assayed enriched transcription factor binding from ENCODE and ChEA datasets using the Enrichr web tool^39–41^. We found *Nfe2l2* (the gene encoding for protein NRF2) consensus sites were the most significantly enriched (**Figure 1D**). NRF2 has previously been implicated in pancreatic tumorigenesis, progression, and metastasis where it regulates expression of genes involved in ROS detoxification, cell metabolism, proliferation, invasion, and migration^20, 22, 42, 43^. We assayed our dataset for NRF2 target genes and found many were significantly modulated in the Cre2 and Cre3 cells compared to GFP controls (**Figure 1E** and **Supplemental Figure 3**). Among the NRF2-target genes with the highest fold change were *Gsta1*, *G6pdx*, and *Atf4*.

Kras-mediated regulation of ROS is complex, being stage, context, and tissue of origin-dependent. In pancreatic cancer, previous work has illustrated that mutant Kras drives Nrf2 to maintain low levels of ROS during tumorigenesis^20^. However, if this ROS is quenched, ADM is blocked^21^. Reciprocally, decreased oxidative PPP activity can promote ROS, which leads to PanIN and metastasis^22^. Our study closely intersects with these studies, uniting the Kras-Nrf2 observation with the oxidative PPP. Specifically, during ADM, we observed upregulation of *G6pdx* and *Pgd*, genes encoding for both rate-limiting oxidative PPP enzymes, glucose-6-phosphate dehydrogenase (G6PD) and 6-phosphogluconate dehydrogenase (PGD), respectively (**Figure 1E**). These enzymes and NADPH-generating cytosolic malic enzyme (ME1) are NRF2 targets. Taken together, these data suggest that upregulated NRF2-related pathways are central in ADM formation.

### G6PD-deficiency decreases oxidative PPP flux in Kras^G12D^-expressing acinar cells

As illustrated in **Figure 2A**, the PPP branches from glycolysis and consists of two arms: oxidative and nonoxidative. According to classical depictions, flux through the oxidative arm generates NADPH and ribose for nucleotides. When there is a demand for NADPH independent of proliferation, excess pentose carbon is recycled through the nonoxidative PPP into the glycolytic intermediates, fructose 6-phosphate and glyceraldehyde 3-phosphate^23^. Contrasting these classical models, in oncogenic *Kras*-driven pancreatic cancer, we previously observed that the increased glycolytic flux led to reverse nonoxidative PPP activity, resulting from overflow metabolism presumably to meet the need for nucleotide biosynthesis^16^.

**Figure 2.**
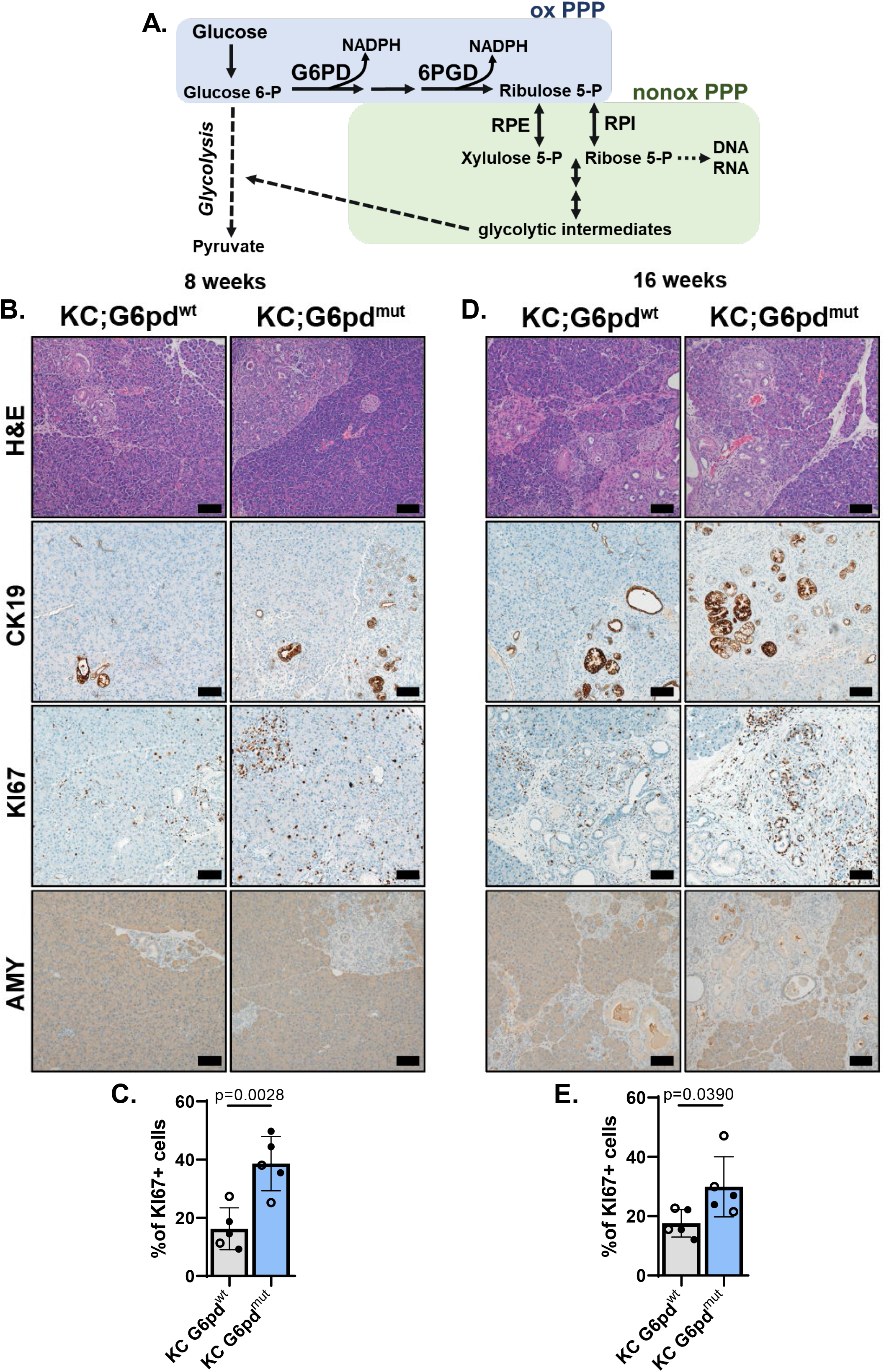
G6PD*-*deficiency accelerates acinar-to-ductal metaplasia in animal model. **A.** Schematic of the pentose phosphate pathway (PPP). In the first step of glycolysis, glucose is converted into glucose 6-phosphate. Glucose-6-phosphate dehydrogenase (G6PD) is the first and rate-limiting enzyme of the oxidative PPP (oxPPP; blue) and converts glucose 6-phosphate (Glucose 6-P) into 6-phosphogluconolactone. During this conversion, G6PD also produces NADPH. 6-Phosphogluconate dehydrogenase (6PGD) is another NADPH-producing enzyme in the oxPPP, which generates ribulose 5-phosphate (Ribulose 5-P). Ribulose 5-P can be used in the nonoxidative PPP (nonoxPPP; green) by enzymes RPE (Ribulose 5-phosphate 3-epimerase) and RPI (Ribose-5-phosphate isomerase) to generate glycolytic intermediates and precursors for nucleic acids. **B.** Hematoxylin & eosin (H&E) staining; cytokeratin 19 (CK19) immunostaining for ductal and ADM cells; KI67 immunostaining for proliferation, and amylase (AMY) immunostaining for acinar cells in pancreas of 8-week-old KC;G6pd^wt^ and KC;G6pd^mut^ mice. Scale bar = 100µm. **C.** Percent of KI67+ cells in pancreas tissue as quantified from male (closed circle) and female (open circle) mice in KC;G6pd^wt^ (grey bar) and KC;G6pd^mut^ (blue bar) mice at 8 weeks. n = 5 for each genotype. **D.** Hematoxylin & eosin (H&E) staining; CK19 immunostaining for ductal, ADM, and early PanIN, KI67 immunostaining for proliferation, and AMY immunostaining for acinar cells in pancreas of 16-week-old KC;G6pd^wt^ and KC;G6pd^mut^ mice. Scale bar = 100µm. **E.** Percent of KI67+ cells in pancreas tissue as quantified from male (closed circle) and female (open circle) mice in KC;G6pd^wt^ (grey bar) and KC;G6pd^mut^ (blue bar) mice at 16 weeks. n = 5 for each genotype.

Thus, we sought to determine the effects of oxidative PPP metabolism on ADM because 1) induction of *Kras^G12D^* mediated a significant increase of the gene coding for G6PD (**Figure 1E**) and 2) this contrasted our previous observation of activation of the nonoxidative PPP enzymes RPE and RPIA^16^. G6PD-deficiency is an X-linked disorder and the most common gene mutation in the world^44^. The oxidative PPP is the only source for NADPH in red blood cells and can cause hemolytic anemia in patients. A mouse model of human G6PD*-* deficiency demonstrates enzyme activity of 15-20% in hemizygous, 50-60% in heterozygous, and 15-20% in homozygous mutant mice^45, 46^. We generated *LSL-Kras^G12D/+^*; *Ptf1a^Cre/+^*; *G6pd^mut^* mice (KC;G6pd^mut^) to determine how G6PD-deficiency, and decreased oxidative PPP flux, alters pancreatic tumorigenesis (breeding scheme in **Supplemental Figure 4**). In these mice, oncogenic *Kras^G12D^* is induced in the exocrine pancreas by *Ptf1a^Cre/+^* (KC)^47^ in a mouse with global G6PD-deficiency.

To determine if oxidative PPP activity was indeed decreased in our KC;G6pd^mut^ mice, we traced radioactive carbon incorporation into carbon dioxide (CO_2_) that is produced from the oxidative PPP and decarboxylation of pyruvate into the TCA cycle (as first described in ^48^). [1-^14^C]glucose labels CO_2_ derived from the oxidative PPP and the TCA cycle; [6-^14^C]glucose labels CO_2_ derived from the TCA cycle and used as a control to determine oxidative PPP activity (**Supplemental Figure 5A**). As expected, KC;G6pd^mut^ acinar cells have decreased oxidative PPP flux, as measured by [1-^14^C]O_2_, compared to mice with wild-type *G6pd* (KC;G6pd^wt^) (**Supplemental Figure 5B**). TCA cycle flux was not affected by *G6pd* status, as measured by [6-^14^C]O_2_ (**Supplemental Figure 5C**). Thus, these data suggest that the KC;G6pd^mut^ acinar cells have less oxidative PPP activity.

### G6PD-deficiency accelerates ADM formation in mouse models

ROS generated during pancreatic tumorigenesis is countered by flux into the oxidative PPP for NADPH and GSH (reduced glutathione) recycling from GSSG (oxidized glutathione)^16, 21, 22, 49, 50^. We hypothesized that decreasing oxidative PPP activity via mutant *G6pd* would promote *Kras^G12D^*-driven ADM and tumorigenesis via increasing oxidative stress. To address this hypothesis, we aged KC;G6pd^mut^ and KC;G6pd^wt^ mice to 8 weeks when ADM is present within the pancreas of KC mice. Indeed, as expected, we observed ADM in KC;G6pd^wt^ mice as noted by histology from H&E-stained tissue sections (**Figure 2B**). We also observed expression of cytokeratin 19 (CK19), increased proliferation (KI67), and decreased amylase (AMY) within ADM lesions. In KC;G6pd^mut^ mice, we observed more ADM lesions and significantly increased proliferation (**Figure 2B, 2C, Supplemental Figure 6A**). Similarly, at 16 weeks of age, when both ADM and early PanIN lesions are present in KC mice, KC;G6pd^mut^ mice developed more ADM and PanIN lesions, marked by an increase in CK19, a decrease in amylase, and significantly more proliferation compared to KC;G6pd^wt^ mice (**Figure 2D, 2E, Supplemental Figure 6A**). While we observed more early lesions in KC;G6pd^mut^ mice when compared to KC;G6pd^wt^ mice, we did not find a difference in pancreas weight to body weight ratios, which is an indirect measure of tissue transformation (**Supplemental Figure 6B**). Our data suggest that G6PD-deficiency, and therefore decreased flux through the oxidative PPP, accelerates early lesion formation in *Kras^G12D^*-driven mouse models.

### G6PD-deficiency accelerates PanIN formation in mouse models

To determine whether G6PD-deficiency influences PanIN formation, we aged mice to 26 weeks, when low grade PanINs (typically PanIN1a/b and occasionally PanIN2) are abundant in KC models^47^. Upon observation of H&E-stained tissue, we noted more transformed area in KC;G6pd^mut^ mice (**Figure 3A**). Additionally, when comparing KC;G6pd^mut^ tissues with KC;G6pd^wt^ tissues, we also noted more CK19 expression and Alcian blue stain (detects mucin production present in PanINs) in KC;G6pd^mut^ pancreata (**Figure 3A, Supplemental Figure 6A**). We did not observe a difference in pancreas weight to body weight ratios in 26-week-old KC;G6pd^mut^ and KC;G6pd^wt^ mice (**Supplemental Figure 6B**). To quantify the extent of tissue transformation, pancreata were histologically graded. KC;G6pd^mut^ tissues contained more PanIN1, PanIN2, and high-grade PanIN3 lesions (**Figure 3B, 3C**). KC;G6pd^mut^ pancreata also showed less acinar cell area and more ADM. Overall, these data suggest decreasing oxidative PPP through mutant *G6pd* accelerates PanIN formation in mice.

**Figure 3.**
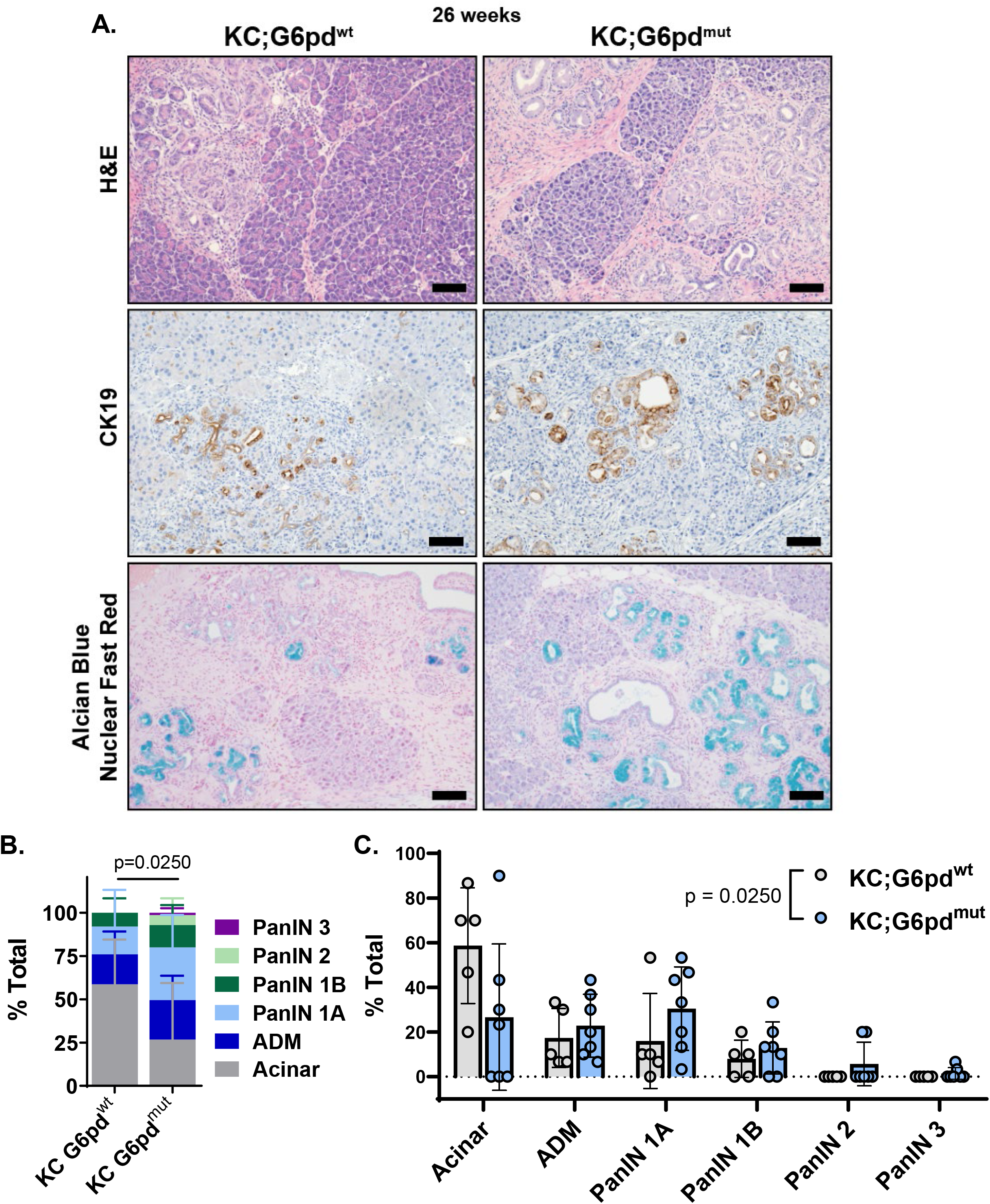
G6PD*-*deficiency accelerates PanIN lesions in animal model of pancreatic tumorigenesis. **A.** Hematoxylin & eosin (H&E) staining; cytokeratin 19 (CK19) immunostaining for ductal and ADM cells; and Alcian blue for PanIN-produced mucin & nuclear fast red counterstain in pancreas of 26-week-old KC;G6pd^wt^ and KC;G6pd^mut^ mice. Scale bar = 100µm. **B.** Pathological grading of KC;G6pd^wt^ and KC;G6pd^mut^ pancreas tissues representing the % total tissue area with acinar cells, ADM, and PanIN lesions. n = 5 KC;G6pd^wt^ mice; n = 7 KC;G6pd^mut^ mice. **C.** Pathological grading of KC;G6pd^wt^ and KC;G6pd^mut^ pancreas tissues from 3B, plotted by Acinar, ADM, and PanIN grades.

### Accelerated ADM in *G6pd*-mutants is due to increased ROS

The oxidative branch of the PPP is a major producer of NADPH, which can reduce oxidized co-factors (e.g. glutathione and thioredoxins) to promote ROS neutralization. Given that ROS is required for ADM formation^21^, we next assessed if there was an increase in ROS in early lesions of *G6pd* mutants. ROS can induce oxidative stress by damaging lipids, proteins, and nucleic acids. One such reactive aldehyde produced by lipid peroxidation is 4-hydroxynonenal (4-HNE). We performed immunostaining for 4-HNE and found higher levels in KC;G6pd^mut^ pancreata compared to KC;G6pd^wt^ (**Figure 4A, 4B**).

**Figure 4.**
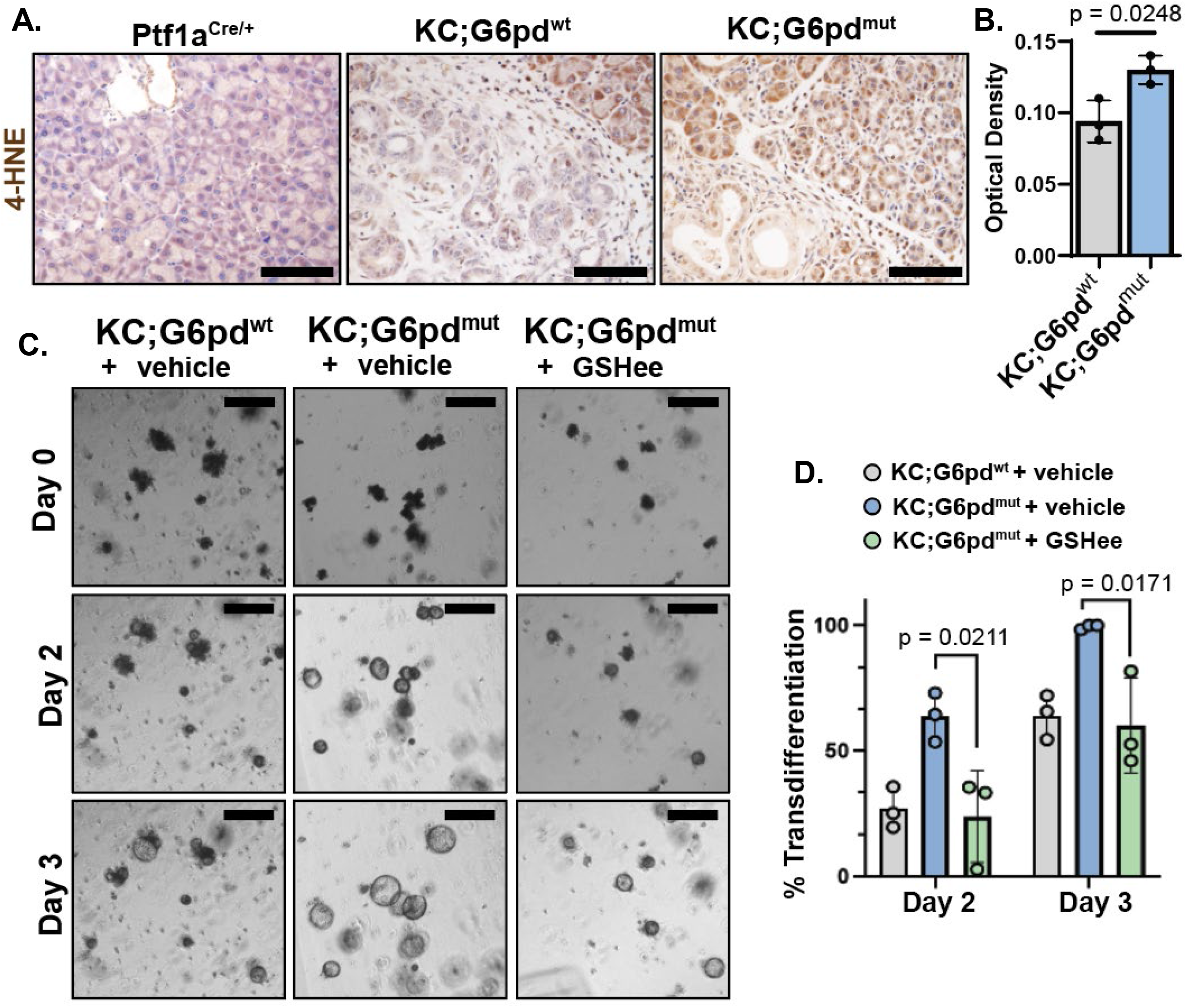
G6PD*-*deficiency increases reactive oxygen species. **A.** Immunostaining for lipid peroxidation marker 4-hydroxynonenal (4-HNE) in 8-week-old Ptf1a^Cre/+^, KC;G6pd^wt^ and KC;G6pd^mut^ pancreas. Scale bar = 100µm. **B.** Quantification of 4-HNE staining. Each datapoint represents one mouse. n = 3 mice per genotype. **C.** Bright field images of *ex vivo* acinar cell cultures from KC;G6pd^wt^ and KC;G6pd^mut^ mice at 2 and 3 days. Cells were treated with mock (vehicle) or 1mM glutathione ethyl ester (GSHee), a cell permeable glutathione. Scale bar = 250µm. **D.** Quantification of acinar cells undergoing ADM in *ex vivo* cultures at 2 days and 3 days. Grey bars are KC;G6pd^wt^ cells mock treated, blue bars are mock-treated KC;G6pd^mut^ cells, green bars are KC;G6pd^wt^ and KC;G6pd^mut^ cells treated with 1mM GSHee. Each datapoint represents one mouse. n = 3 mice per genotype.

We hypothesized that accelerated ADM formation with *G6pd-*deficiency could be rescued by the addition of antioxidants to lower ROS levels. To test this, we isolated acinar cells from KC;G6pd^mut^ and KC;G6pd^wt^ mice, 3D-embedded the cells in Matrigel, and cultured the cells in limited media^12^. We imaged cells for 3 days and measured acinar to ductal cyst conversion, or the percent of cells that have transdifferentiated to ADM. Similar to our result *in vivo,* KC;G6pd^mut^ cells undergo more rapid transdifferentiation in *ex vivo* culture compared to KC;G6pd^wt^ mice (**Figure 4C, 4D**). Over 50% of KC;G6pd^mut^ cells transdifferentiate into ductal-like cysts at Day 2, compared to less than 25% of KC;G6pd^wt^ cells at the same timepoint (**Figure 4D**). The increased ADM formation in KC;G6pd^mut^ cells is rescued with a cell permeable glutathione, glutathione ethyl ester (GSHee) (**Figure 4C, 4D**). These data suggest that KC;G6pd^mut^ acinar cells have increased ROS levels, which promotes faster progression of early lesions.

### G6PD-deficiency does not affect survival in PDAC mouse models

We next questioned if G6PD-deficiency plays a role in pancreatic cancer progression and survival. Since most KC mice do not develop invasive carcinoma^47^, we used a KPC mouse model (*LSL-Kras^G12D/+^*; *LSL-Tp53^R172H/+^*; *Ptf1a^Cre/+^*) to include additional loss of tumor suppressor Trp53, which increases tumorigenesis, promotes invasive carcinoma and metastasis, and decreases survival^51^. We generated KPC mice with G6PD-deficiency to determine if accelerated tumorigenesis enhances invasive PDAC and decreases survival. We observed no difference in pancreas weight to body weight ratios between KPC;G6pd^mut^ and KPC;G6pd^wt^ mice (**Figure 5A**). Advanced disease was present in both KPC;G6pd^mut^ and KPC;G6pd^wt^ mice (**Figure 5B**), but there was no statistically significant difference in survival (**Figure 5C**). While G6PD-deficiency promotes precancerous lesions in KC mice, these data indicate that G6PD-deficiency does not affect survival in KPC mice.

**Figure 5.**
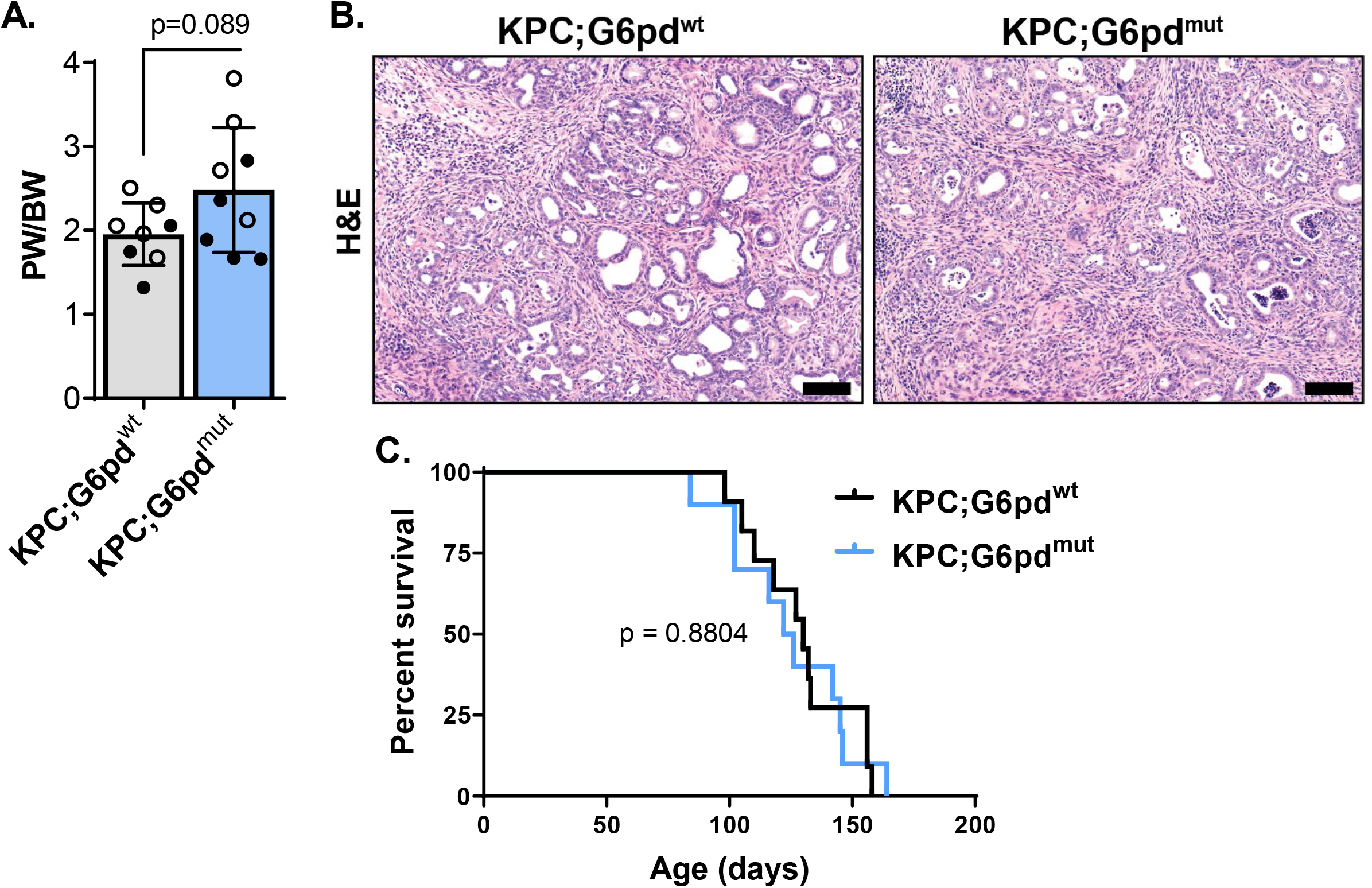
G6PD*-*deficiency does not decrease overall survival in KPC mice. **A.** Pancreas weight (PW) to body weight (BW) ratios in 90-day-old male (closed circle) and female (open circle) KPC;G6pd^wt^ and KPC;G6pd^mut^ mice. Each datapoint represents one mouse. n = 8 KPC;G6pd^wt^ mice; n = 9 KPC;G6pd^mut^ mice. **B.** Hematoxylin & eosin (H&E) staining in 90-day-old KPC;G6pd^wt^ and KPC;G6pd^mut^ pancreas. Scale bar = 100µm. **C.** Kaplan–Meier survival curve of KPC;G6pd^wt^ and KPC;G6pd^mut^ mice. n = 11 KPC;G6pd^wt^ mice; n = 10 KPC;G6pd^mut^ mice.

## DISCUSSION

Basal ROS levels are elevated in malignant cells from increased metabolism, which can potentiate mitogenic signaling. However, antioxidant programs must be initiated to restrain ROS from reaching cytotoxic levels. *Kras^G12D^* elevates both ROS production and antioxidant pathways during ADM and pancreatic tumorigenesis^20,21^. We and others show that antioxidant treatment impedes *ex vivo* ADM formation and tumorigenesis *in vivo* by lowering ROS levels^21, 52^. In contrast, abrogating antioxidant responses through genetic deletion of NRF2, a primary antioxidant regulator, causes exceedingly high levels of ROS that induce cellular senescence^20^. Collectively these results illustrate of the delicate balance of ROS levels in tumorigenesis.

We similarly observed an NRF2 signature in our *ex vivo* ADM culture model, suggesting that oncogenic *Kras* instructs NRF2-dependent ROS control in precancerous lesions. An important effector of the antioxidant response during pancreatic tumorigenesis is TP53-induced glycolysis and apoptosis regulator (TIGAR), which dampens ROS by redirecting glycolytic flux into the oxidative PPP for NADPH generation^22^. We inhibited oxidative PPP flux through mutation of *G6pd*, the first and rate limiting step of the oxidative PPP. G6PD-deficiency increased the rate of ADM in *ex vivo* acinar cultures and tumorigenesis in KC mice. Our data suggests that this acceleration is due to increased oxidative stress from decreased production of NADPH reducing equivalents. However, we saw no change in survival with G6PD-deficiency in KPC mice, pointing to clear differences in NADPH and ROS control in precancerous lesions compared to advanced disease. A similar, yet distinct, phenomena has been described with the KEAP1/NRF2 pathway in lung tumorigenesis, where NRF2 activation (by *Keap1* deletion) promotes oncogenic *Kras*-induced tumor initiation but impairs tumor progression^53^.

We have shown that PDAC cells display less dependence on the oxidative PPP and instead rewire glutamine metabolism to generate NADPH and uncouple the nonoxidative PPP branch to generate nucleotide precursors^16, 17^. Therefore, in disease progression, G6PD may no longer be important for NADPH production once glutamine metabolism is reprogrammed to produce NADPH. PDAC cells may bypass the need for G6PD-dependent NADPH production with G6PD-deficiency by instead relying on other NADPH-producing enzymes, like ME1 and IDH1, as observed in melanoma^54^, lung cancer^55^, and colon cancer^56^ models. These results provide additional evidence and context for divergent roles of ROS, redox, and antioxidant pathways in tumor initiation, development, progression, and metastasis.

Pancreatic cancer is among the most lethal cancers^1, 2^. There is a need for early disease detection methods and more effective therapies to improve the dismal prognosis patients face. Historically, KRAS has proven notoriously difficult to target pharmacologically^57^. However, the FDA has approved a KRAS^G12C^ inhibitor, Sotorasib for lung cancer, and Kras^G12C^ inhibitors Sotorasib and Adagrasib show promising results in PDAC clinical trials^58, 59^. Unfortunately, G12C mutations are only found in ∼2% of PDAC patients^60^. Recently, a small molecule KRAS^G12D^ inhibitor, MRTX1133, led to tumor regression in autochthonous PDAC mouse models with an intact immune system^61^. Additionally, the development of a pan-KRAS selective inhibitor may have broad therapeutic implications in PDAC and other KRAS-driven cancers^62^. Bringing our study into the greater perspective, an understanding of the metabolic pathways that initiate PDAC could reveal vulnerabilities in established tumors that can be exploited for treatment or reveal biomarkers for diagnosis of early disease.

## ACKNOWLEDGEMENTS

We thank the members of the Lyssiotis, Crawford, and Shah laboratories and the Pancreatic Disease Initiative at the Rogel Cancer Center, University of Michigan for their valuable comments and input.

## SUPPORT

M.D.R. was supported by NIH grants T32HD007505 and F32CA275283. B.S.N. was supported by NIH grants T32DK094775 and T32CA009676. C.J.H. was supported by a National Pancreas Foundation Research Grant, a Sky Foundation Research Grant, a V Scholar Award (V2021-026), the Chao Family Comprehensive Cancer Center support grant P30CA062203, and NIH grants F32CA228328, K99/R00CA241357, and P30DK034933. J.A.C. was supported by NIH grants R01ES028802, R21CA273646, and P30ES017885. Y.M.S was supported by NIH grants R01CA148828, R01CA245546, and R01DK095201. C.A.L. was supported by the NCI (R01CA248160), a Pancreatic Cancer Action Network/AACR Pathway to Leadership award (13-70-25-LYSS); Junior Scholar Award from The V Foundation for Cancer Research (V2016-009); Kimmel Scholar Award from the Sidney Kimmel Foundation for Cancer Research (SKF-16-005); and a 2017 AACR NextGen Grant for Transformative Cancer Research (17-20-01-LYSS). The funders had no role in study design, data collection and analysis, decision to publish or preparation of this manuscript.

## DISCLOSURES

In the past three years, C.A.L. has consulted for Astellas Pharmaceuticals, Odyssey Therapeutics, Third Rock Ventures, and T-Knife Therapeutics, and is an inventor on patents pertaining to Kras regulated metabolic pathways, redox control pathways in pancreatic cancer, and targeting the GOT1-ME1 pathway as a therapeutic approach (US Patent No: 2015126580-A1, 05/07/2015; US Patent No: 20190136238, 05/09/2019; International Patent No: WO2013177426-A2, 04/23/2015). G.C. was scientific founder of Decibel Therapeutics, had equity interest in the company, and received compensation for consulting. The company was not involved in this study.

## MATERIALS AND METHODS

### Mouse strains

*LSL-Kras^G12D/+^* mice were maintained on a C57BL/6 background. G6PD-deficient mice have been previously described^44–46^. *LSL-Kras^G12D/+^*; *Ptf1a^Cre/+^*; *G6pd^mut^*or *G6pd^wt^* mice were maintained on a mixed background. *LSL-Kras^G12D/+^*; *LSL-Tp53^R172H/+^*; *Ptf1a^Cre/+^*; *G6pd^mut^* or *G6pd^wt^* mice were maintained on a mixed background.

### Genotyping

In-house genotyping was done using DNA extracted from tail or ear tissue using DirectPCR Lysis Reagent (Viagen Biotech, Inc). Thermal cycling was performed using the Power SYBR Green 2X Master Mix (ThermoFisher). Gel electrophoresis was performed on 2.5% agarose gel with 10,000X SYBR Safe DNA gel stain (Invitrogen). 1 kb and 100 bp DNA Ladders (New England Biolabs) were used to measure amplicon size. Gels were run at 125V for 75 minutes and imaged using the ChemiDoc Imaging System (Bio-Rad). The primers utilized for in-house PCR were as follows: Cre (For-TCGCGATTATCTTCTATATCTTCAG, Rev-GCTCGACCAGTTTAGTTACCC); LSL-Kras^G12D^ (Universal-CGCAGACTGTAGAGCAGCG, Mutant-CCATGGCTTGAGTAAGTCTGC). LSL-Trp53^R172H^ (p53-Wildtype-TTACACATCCAGCCTCTGTGG, p53-5’-AGCTAGCCACCATGGCTTGAGTAAGTCT, p53-3’-CTTGGAGACATAGCCACACTG). G6pd mouse PCR primers were made between exon 1 and intron 1, around the reported mutation site (Nicol 2000) (sense: GGAAACTGGCTGTGCGCTAC, antisense: TCAGCTCCGGCTCTCTTCTG). The PCR product was then digested using DdeI restriction enzyme and run on a 3% agarose gel. *G6pd wildtype* have two bands (214 and 55bp), *G6pd mutant homozygotes* have 1 band (269bp), *G6pd mutant heterozygotes* have 3 bands (269, 214, and 55bp).

Following the global pandemic, genotyping was outsourced to a commercial vendor, who used real time PCR with specific probes designed for each gene (Transnetyx, Cordova, TN).

### Animal use

Mouse experiments were conducted under the guidelines of the Office of Laboratory Animal Welfare and approved by the Institutional Animal Care and Use Committees (IACUC) at the University of Michigan under protocol PRO00010606. All mice were kept in ventilated racks in a pathogen-free animal facility with a 12 h light/12 h dark cycle, 30–70% humidity, and 68–74°F temperatures maintained in the animal facility at Rogel Cancer, University of Michigan. Mice were overseen by the Unit for Laboratory Animal Medicine (ULAM). Male and female mice were used for each experiment.

### Three-dimensional primary acinar cell culture

Pancreas from mice aged 10 weeks was harvested and rinsed twice in 5 mL cold HBSS (Gibco). Tissue was minced with sterile scissors into 1-5 mm sized pieces then centrifuged for 2 minutes at 300 g and 4°C. Media was aspirated and minced tissue was digested with ∼5 mg of Collagenase P (Roche) in 5 mL cold HBSS for 15-20 minutes, shaking at 100 rpm at 37°C. Collagenase P was inhibited by addition of 5 mL cold 5% fetal bovine serum in HBSS. Cells were centrifuged for 2 minutes at 300 g and 4°C then washed with 5 mL cold 5% FBS in HBSS. This was repeated twice more. Cells were passed through 500 μm strainer (Pluriselect, Fisher, NC0822591), then through a 100 μm polypropylene mesh (Fisherbrand, 22363549), and then pelleted through 10 mL of 30% fetal bovine serum gradient. Cells were resuspended in media and incubated at 37°C for at least 4 hours prior to plating. Cells were cultured in 1x Waymouth’s media (11220035, Gibco) supplemented with 0.4 mg/mL soybean trypsin inhibitor (Gibco, 17075-029), 1 ug/mL dexamethasone (Sigma, D4902), and 0.5% gentamicin (Lonza, 17-519L). All media was sterilized through a 0.22 μm PVDF membrane (Millipore, Stericup Filter Unit, SCGVU01RE; Steriflip Filter Unit, SE1M179M6). Acinar cells were grown free floating in 6-well plates (for RNA isolation) or embedded in Matrigel (Corning, 354234) and plated in 24-well plates for measuring transdifferentiation rates. Culture media was added on top of solidified matrix and changed on days 1, 3, and 5 after plating. Cells were treated with water (vehicle) or 1mM glutathione ethyl ester (GSHee; Cayman Chemical, 14953) dissolved in DMSO. Cells were imaged using EVOS FL Auto Imaging System (ThermoFisher Scientific).

### Adenovirus treatment in primary acinar cell cultures

Acinar cells were isolated from *LSL-Kras^G12D/+^* mice and infected with either control adenovirus (ad-GFP) or adenovirus that expresses Cre recombinase (ad-CRE) to induce expression of mutant Kras^G12D^. RNA was isolated 24 hours after plating ad-GFP infected acinar cells (GFP) and 24, 48, and 72 hours after plating ad-CRE infected cells (Cre1/2/3, respectively).

### RNA isolation and purification from primary acinar cell cultures

Primary acinar cell cultures were harvested by centrifugation for 2 minutes at 300 g. The pellet was washed once with ice cold DPBS and lysed in RLT+ buffer containing 1% β-mercaptoethanol (Sigma, M6250), which was then passed through a Qiashredder column (Qiagen, 79654). RNA was purified using a RNeasy Plus Mini Kit (Qiagen, 74136) and analyzed on a Nanodrop 2000c (Thermo Scientific) spectrophotometer for quantification. Purity was assessed based on the A260/A280 ratio. RNA quality was further assessed using a total RNA kit on a 2100 Bioanalyzer (Agilent Technologies) and using the generated RIN number to determine quality, and all samples sequenced had a RIN greater than 7.5. 1µg of each sample was submitted to the Mayo Clinic Genome Analysis Core using the Illumina HiSeq 2000 platform and Illumina TruSeq v2 mRNA library prep. Data were generated and processed through the Mayo Clinic Genome Analysis Core pipelines.

### RNA-sequencing data analysis

RNA-sequencing counts matrices were analyzed in edgeR to identify differentially expressed genes between experimental conditions. The expression data were filtered to remove genes with low counts using the edgeR filterByExpr() function with default settings. Normalization factors and dispersion parameters were then calculated prior to generating a log2-transformed counts per million (cpm) matrix for analysis. Overall gene expression patterns per sample were first visualized with a multidimensional scaling plot. Differential gene expression between each timepoint (Cre1/2/3) and GFP control was estimated using the quasi-likelihood negative binomial generalized log-linear approach in edgeR. Genes were considered differentially expressed between a treatment and control at a false discovery rate (FDR) adjusted p-value less than 0.05. Enriched differentially expressed genes per treatment were identified using the topGO() and topKEGG() functions in edgeR. Enriched gene sets for the entire list of differentially expressed genes per treatment were identified using Gene Set Enrichment Analysis software. To identify enriched transcription factor binding in differentially expressed gene lists, the top 500 overexpressed genes within a condition were uploaded to the Enrichr web tool^39–41^ to quantify enrichment for ENCODE and ChEA consensus transcription factor targets. We visualized gene expression patterns for key markers of acinar and ductal cell types as well as NRF2 transcription factor targets using pheatmap() in R. Analyses were conducted using R software version 4.2.2.

### ^14^C glucose incorporation into CO_2_

Cells were treated with 1 μCi 1-^14^C (Perkin Elmer, NEC043X050UC) or 6-^14^C glucose (Perkin Elmer, NEC045X050UC) and incubated at 37°C for 4 hours. To release ^14^CO_2_, 150 µL of 3 M perchloric acid (Sigma, 244252) was added to each well and immediately covered with phenylethylamine (Sigma, P6513)-saturated Whatman paper and incubated at room temperature overnight. The Whatman paper was then analyzed by scintillation counting (Beckman, LS6500) and normalized to surrogate protein quantification.

### Tissue processing for histology

Mice were euthanized by CO_2_ asphyxiation or isoflurane overdose, followed by cervical dislocation. The tissue was quickly harvested and fixed overnight at room temperature with zinc formalin fixative (Z-Fix; Anatech, Ltd and Cancer Diagnostics, Inc). Fixed tissues were then transferred to 70% ethanol. Tissues were processed using a Leica ASP300S Tissue Processor (Leica Microsystems, Inc), paraffin embedded, and cut into 5 μm sections.

### Immunohistochemistry

Tissue sections were deparaffinized with Histo-Clear (National Diagnostics), and re-hydrated with graded ethanol and water. Samples were quenched with 1.5% hydrogen peroxide in 100% methanol for 15 minutes at room temperature. Antigen retrieval was performed in sodium citrate buffer (2.94 g sodium citrate, 500 μl Tween 20, pH 6.0) using a pressure cooker on high or by maintaining slides at a rolling boil for 20 minutes. Slides were blocked in 2.5% bovine serum albumin (Sigma), 0.2% Triton X-100, in PBS for 45 minutes to 1 hour at room temperature. Slides were incubated overnight at 4°C in primary antibodies. After rinsing with PBS, slides were incubated for 1 hour at room temperature in secondary antibodies. Blocking of endogenous biotin, biotin receptors, and avidin binding sites was performed using Avidin/Biotin Blocking Kit (Vector Laboratories) per manufacturer’s protocol. Substrate reaction and detection was performed using DAB Peroxidase (HRP) Substrate Kit (With Nickel), 3,3’-Diaminobenzidine (SK4100, Vector Laboratories) as detailed per the manufacturer’s protocol. Slides were counterstained, mounted in Permount Mounting Medium (Fisher Scientific), coverslipped, and allowed to dry overnight before imaging.

### Counter stains and special stains

Hematoxylin and eosin (H&E) staining was performed using Mayer’s hematoxylin solution and Eosin Y (Thermo Fisher Scientific). Blue and Nuclear Fast Red staining was performed using Alcian Blue (pH 2.5) Stain Kit (H-3501, Vector Laboratories) per manufacturer’s protocol.

### Antibodies

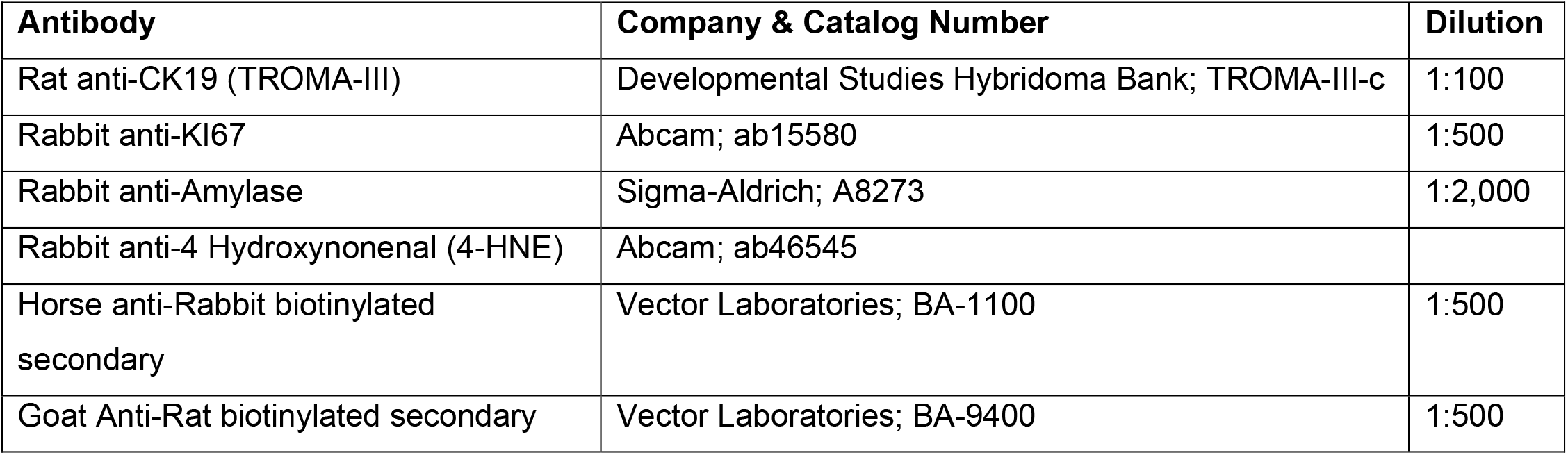

### Imaging and Quantifying Staining

Slides were imaged with CellSens Standard software using an Olympus BX53F microscope, fitted with an Olympus DP80 digital camera (Olympus Life Science). To quantify staining, 3-5 images were taken per slide at 10x magnification. Images were anonymized, randomized, and positive DAB signal was quantified using QuPath software^63^. For tissue grading, 3 images were taken per slide at 20x magnification in a blinded manner and graded by a pathologist in a blinded manner.

### Statistical analysis

Statistics were calculated using GraphPad Prism 8 and analyzed with a Student’s t-test (unpaired, two-tailed) when comparing two groups to each other, one-way ANOVA analysis with Tukey post hoc test with groups greater than 2 with a single variable, and two-way ANOVA with Tukey post hoc test with groups greater than two multiple variables. All data are presented as mean ± standard deviation.

---

**Supplemental Figure 1.**
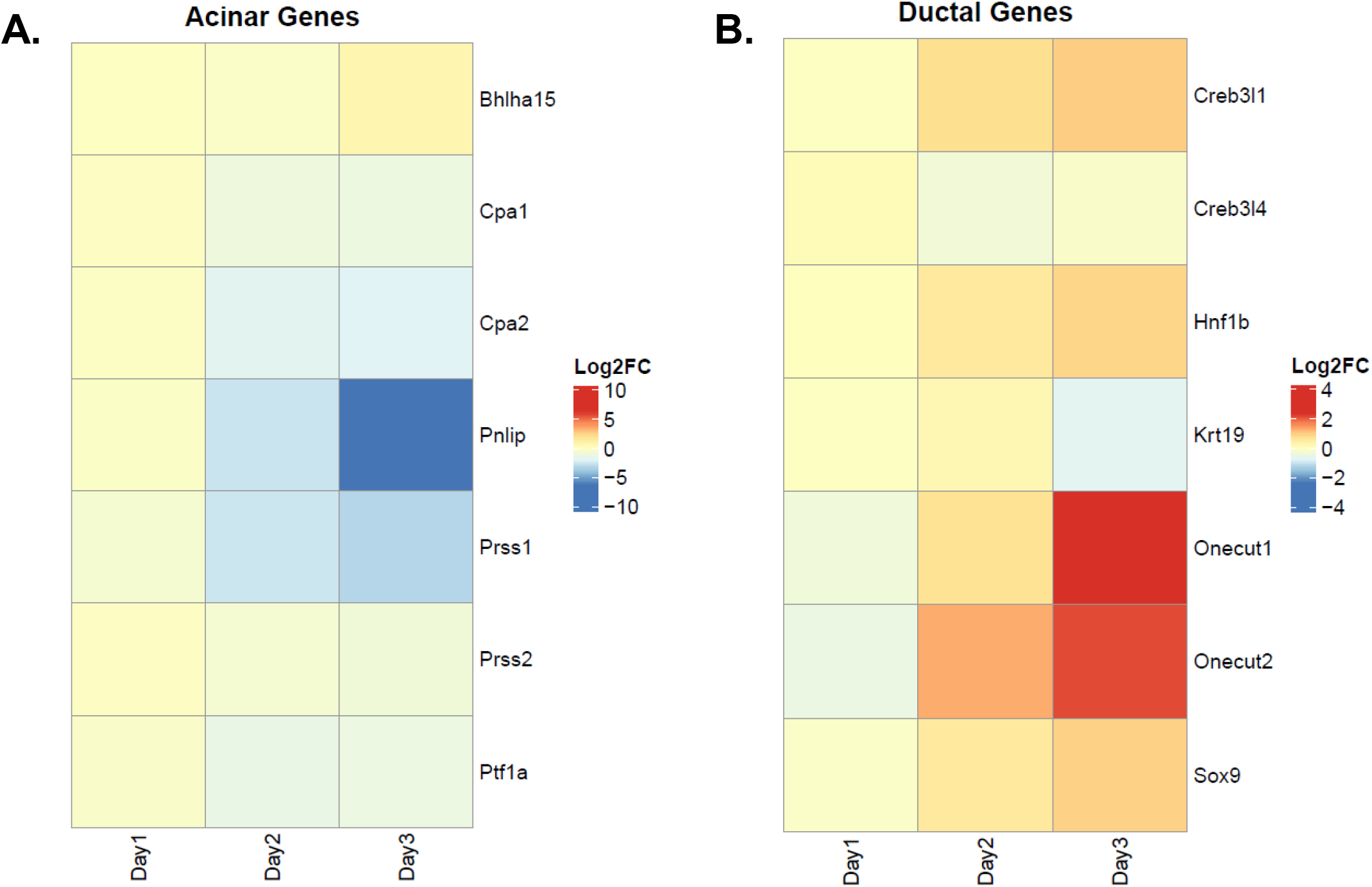
Heatmaps of acinar and ductal genes from RNA-sequencing. **A.** Log_2_FC expression of several acinar genes from RNA-sequencing of *ex vivo* acinar cell cultures at Day 1, 2, or 3 compared to GFP. **B.** Log_2_FC expression of several ductal/ADM genes from RNA-sequencing of *ex vivo* acinar cell cultures at Day 1, 2, or 3 compared to GFP.

**Supplemental Figure 2.**
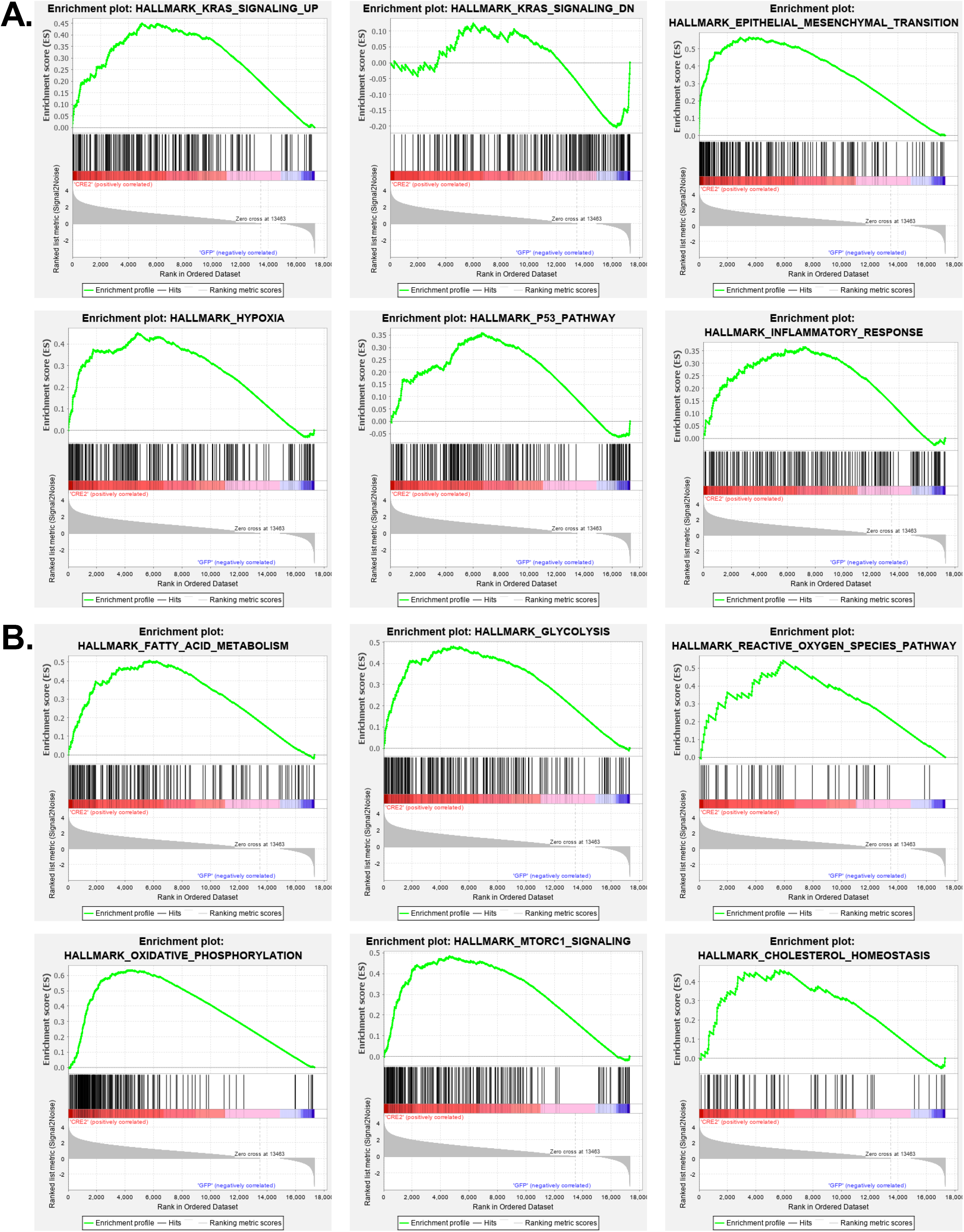
Differentially expressed pathways and GSEAs from RNA-sequencing. **A.** Selected Hallmark GSEA plots of known pathways in ADM that are enriched in *Kras^G12D^*-expressing acinar cells (Cre2 vs. GFP). **B.** Selected Hallmark GSEA plots of metabolic pathways that are enriched in *Kras^G12D^*-expressing acinar cells (Cre2 vs. GFP). Enrichment score (ES) signifies the degree a gene set is overrepresented at the top or bottom of a ranked list of genes. The black vertical bars show where genes within the gene set appear in ranked list. The waterfall plot represents a gene’s correlation with a phenotype.

**Supplemental Figure 3.**
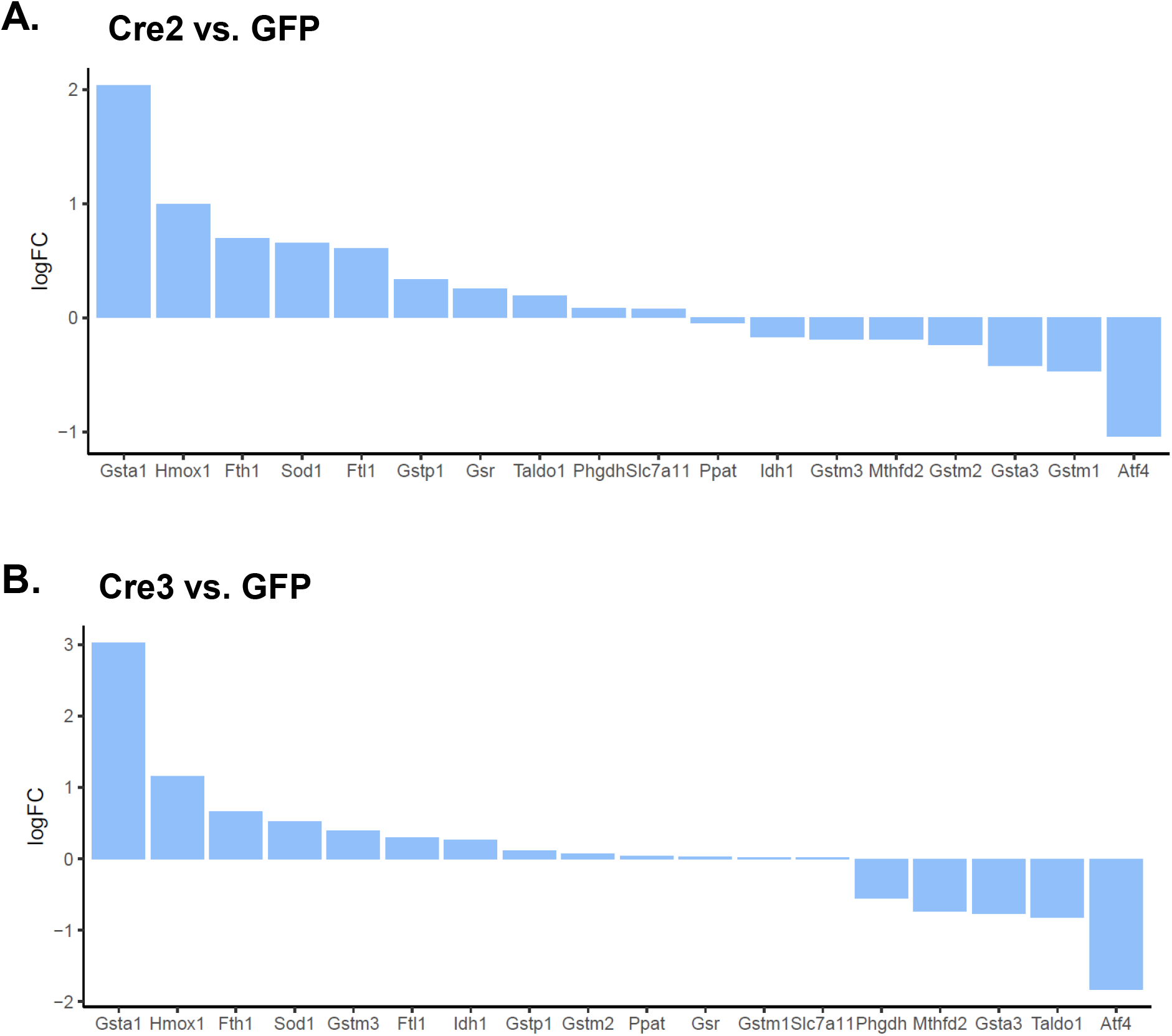
Additional NRF2 target genes from RNA-sequencing. **A.** LogFC expression of additional NRF2-target genes from RNA-sequencing of *Kras^G12D^*-expressing acinar cells when comparing Cre2 to GFP. **B.** LogFC expression of additional NRF2-target genes from RNA-sequencing of *Kras^G12D^*-expressing acinar cells when comparing Cre3 to GFP.

**Supplemental Figure 4.**
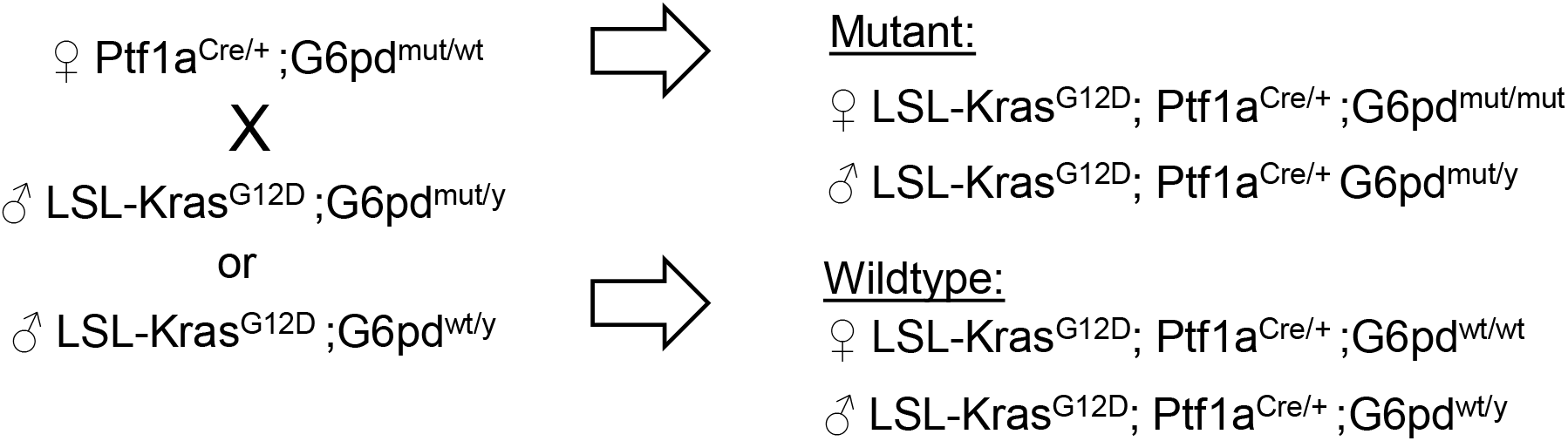
Breeding scheme to generate KC;G6pd^mut^ and KC;G6pd^wt^ mice. Mice with mutant *G6pd*, that mimics human G6PD-deficiency, were bred into the KC (*LSL-Kras^G12D/+^; Ptf1aCre*) line. *Ptf1aCre* was always maintained in female breeders. Experiments used both male and female *G6pd* mutant mice, where females were homozygous for mutant *G6pd* (*G6pd^mut/mut^*) and males were hemizygous for mutant *G6pd* (*G6pd^mut/y^*), as *G6pd* is an X-linked gene. *G6pd* wildtype mice used in experiments were obtained in the same colony and age matched littermates of *G6pd* mutant mice when possible. Female *G6pd* wildtype mice have two wildtype copies of *G6pd* (*G6pd^wt/wt^*) and males have one wildtype copy (their only copy) of *G6pd* (*G6pd^wt/y^*). In the schematics and labelling, “y” in male mice refers to the y chromosome, which does not contain a copy of *G6pd*.

**Supplemental Figure 5.**
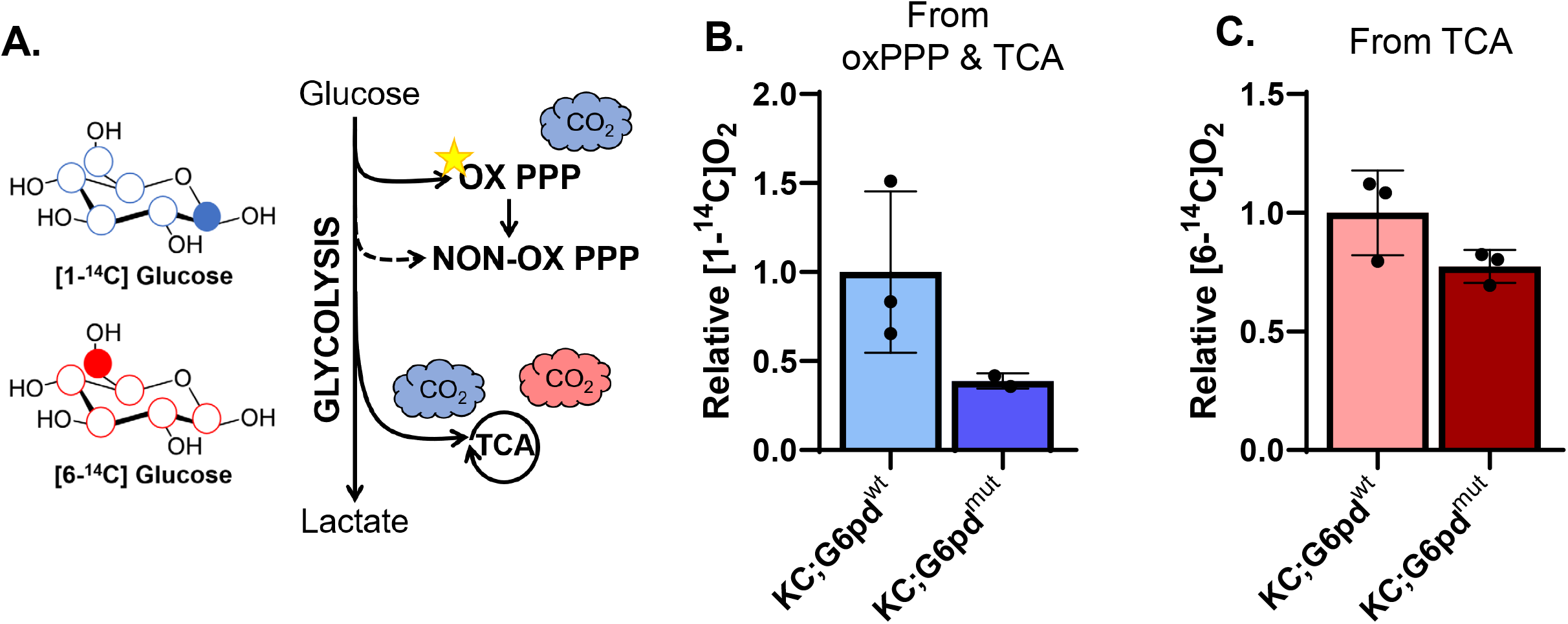
1^4^C-labeling in acinar cells. **A.** Schematic of ^14^C-labeling experiment. [1-^14^C]glucose (blue) can be used in both the oxidative pentose phosphate pathway (ox PPP) and the TCA cycle. [6-^14^C]glucose (red) can be used in the TCA cycle. **B.** Relative amount of [^14^C]-labeled CO_2_ derived from [1-^14^C]glucose. [1-^14^C]O_2_ is generated from either the oxidative pentose phosphate pathway or the TCA cycle. **C.** Relative amount of [^14^C]-labeled CO_2_ derived from [6-^14^C]glucose. [6-^14^C]O_2_ is only generated from the TCA cycle. Each point in B and C represents technical replicates from one mouse.

**Supplemental Figure 6.**
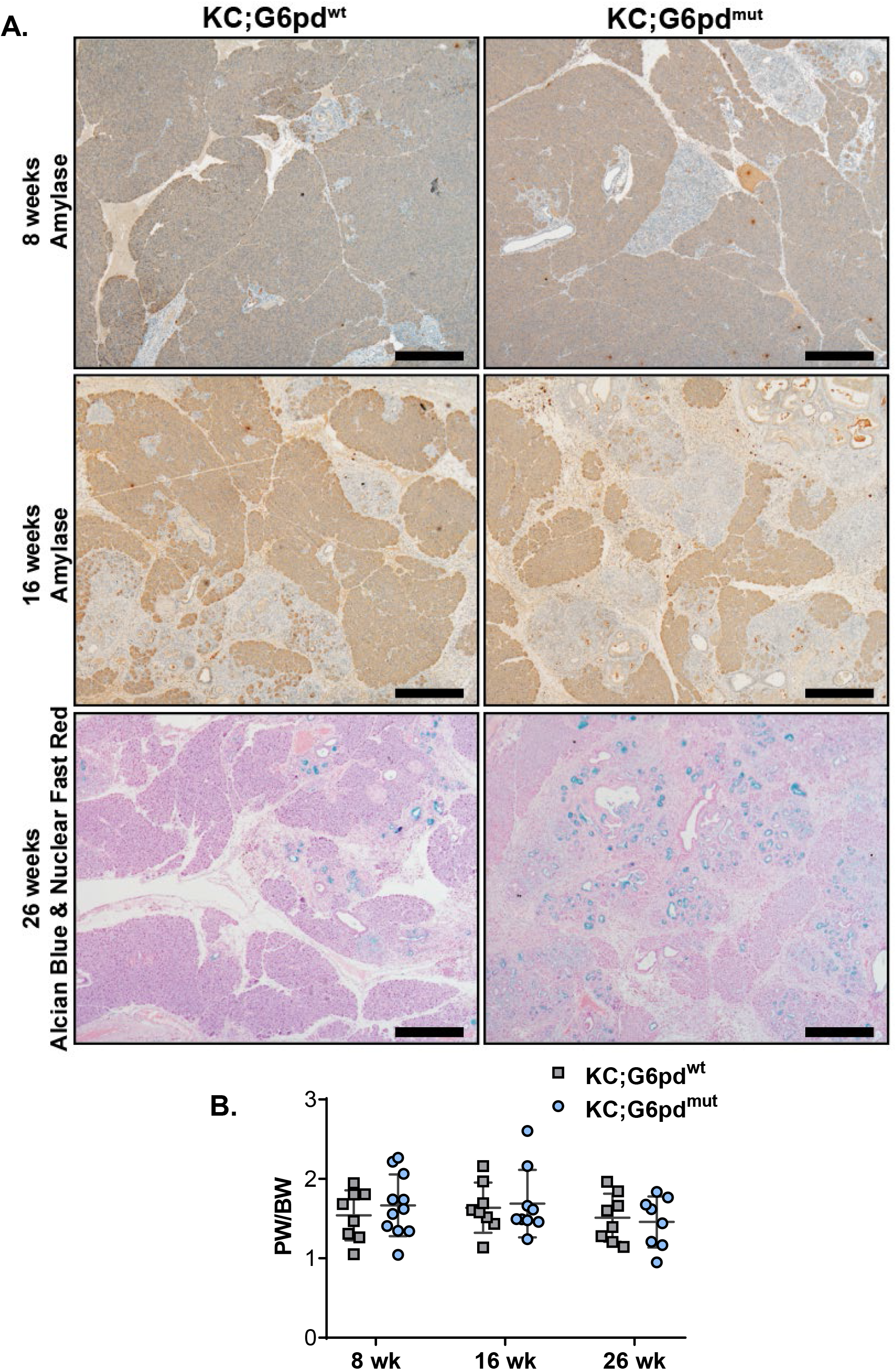
Low magnification images of Amylase and Alcian blue in pancreata and pancreas weight to body weight ratios. **A.** Immunostaining for Amylase (AMY), in 8-week and 16-week-old KC;G6pd^wt^ and KC;G6pd^mut^ pancreas. Alcian blue (PanIN-produced mucin) & nuclear fast red counterstain in pancreas of 26-week-old KC;G6pd^wt^ and KC;G6pd^mut^ mice. Scale bar = 500µm. **B.** Pancreas weight (PW) to body weight (BW) ratios in 8-, 16-, and 26-week-old KC;G6pd^wt^ and KC;G6pd^mut^ mice.

